# How simple physics drives the earliest stages of embryogenesis

**DOI:** 10.64898/2026.01.07.698115

**Authors:** Alaina Cockerell, Peyman Shadmani, Krasimira Tsaneva-Atanasova, David M. Richards

## Abstract

The initial stages of mammalian embryo development involve a single fertilised egg that repeatedly divides to create a solid ball of cells called a morula. Despite the apparent simplicity of this process, which involves only one cell type and a few tens of cells, there are still a host of unanswered questions, particularly around the underlying biophysical mechanisms that are at play. To address this, we here develop a novel type of vertex model that includes cortical tension, cell-to-cell adhesion, membrane curvature and cell volume forces, along with a zona pellucida, cell division and the effect of noise. We fit our model to both mouse and human experimental data, which allows us to address a number of key questions including the relative roles of adhesion and tension, how the cortical tension varies around the cell, the purpose of the zona pellucida, and the rules governing the first few cell divisions. We also determine the biophysical effects responsible for compaction and internalisation, including addressing why the morula does not typically decompact during internalisation. Next, we investigate the position-versus-polarisation debate during trophectoderm differentiation, how the division axis is determined during later divisions, and the role of noise. Finally, we compare human and mouse, focussing on the key similarities that may span all mammals. Our use of a force-based computational model allows us to address fundamental questions relating to mammalian development, particularly the underlying biophysical rules governing early embryogenesis, with important applications to stem cell models such as blastoids, conservation efforts of endangered species and embryo grading during IVF.

**Significance Statement:** The very first stages of embryogenesis are a vital but still poorly understood part of development, with important applications to fertility, fertility lifespan and assisted conception such as IVF. Most research in this area has focussed on experimental approaches, ignoring the potential for biophysical modelling. Here, we address this by developing a novel computer simulation of the first thee-to-four cell divisions of the fertilised egg, resulting in a compact bunch of cells called a morula. In particular, we develop a new type of vertex model for the early embryo that for the first time includes contributions from a range of realistic biophysical forces. By analysing real mouse and human embryo data through our model, we reveal how simple physics drives crucial early developmental processes, including compaction, internalisation, cleavage, noise and species-specific differences.

## 1. Introduction

Development of multicellular organisms is a highly regulated process, with the precise spatial arrangements of different cell types along with precise timing of each stage necessary to achieve viable and successful mature organisms (1, 2). Often the developmental programmes across mammalian species are remarkably similar, governed by conserved biological and biophysical processes (2–5).

This is especially the case during the very earliest stages of embryogenesis, where a single fertilised egg (*i*.*e*. a zygote) undergoes its first few divisions. These divisions of the first handful of cells (called blastomeres) are normally cleavage divisions, where cells divide in two without any significant subsequent growth. This means that the overall embryo volume is roughly conserved over the first rounds of division, forming a multicell solid mass of blastomeres called a morula. The number of cells in the mature morula varies between organism, but is typically around 16-32, taking about two days to form in mouse and three in human.

In most mammals, the morula undergoes compaction, where the contact area between cells expands. This is followed by internalisation, during which a subset of cells move to the middle of the morula and no longer have any contact with the outside medium. Cells that remain on the outside of the embryo are fated to become trophectoderm, whereas those inside become the inner cell mass (ICM) (2, 6–8).

There are a host of open questions related to these earliest developmental stages, including the relative roles of cell-to-cell adhesion and cortical tension, the dynamics of the actin cytoskeleton (9, 10) and how the cell division axes are determined. There is also an ongoing debate about how outer cells become trophectoderm: is it because they are exposed to the outside surface or because they inherit an apical domain during division (11–13)? In addition, the similarities and differences between organisms, such as between human and mouse, is a fascinating and poorly studied area, with relevance to a multitude of fundamental biological and biophysical areas.

Mathematical and computational models have been extensively used across biology and medicine, transforming how we decode life’s complexities in health and disease. Intimately combining modelling approaches with experiments and real data can provide understanding and advances that could not otherwise be achieved (14). For example, models can help predict and identify what is missing from current understanding (15) and can be used to study questions that cannot be tackled experimentally (such as removing cell-to-cell adhesion without any knock-on effect to signalling pathways) (16). At its heart, modelling attempts to simplify a system and abstract the most important components, with the aim of making testable predictions, streamlining experiments and reducing the research cost and time. Perhaps one of the most important consequences of this is that mathematical and computational approaches are often able to identify underlying concepts that span systems, species and even disciplines, providing knowledge that would otherwise be next to impossible to obtain.

There have been a number of previous models of morula formation. Some have focussed on the effect of the zona pellucida (ZP), the extracellular-matrix shell that surrounds the embryo, often using a particle-potential type of model. For example, Krupinski et al. looked at the orientation of the first two cells compared to the long axis of an ellipsoidal ZP, finding that less than a 1% difference in the ZP axis lengths is sufficient to position cells along the long axis (17). In contrast, Giammona et al. focused on the role of the ZP in the packing configurations of the first four cells across several species, identifying that the ZP greatly decreases packing degeneracy and aids correct positioning of early blastomeres (18).

Compaction was first modelled by Goel et al. using a vertex model and energy minimisation (19). This was subsequently expanded by Le Guillou et al. and Maître et al. using a 3D cellular Potts model and a 3D multi-material mesh-based surface-tracking method respectively (9, 20–23). They discovered that increasing adhesion between cells was sufficient to induce compaction and that compaction could be achieved by changing the surface tensions at the cell-medium interface and the cell-to-cell boundary. Maître et al. also used the same framework to investigate internalisation, finding that differences in the cell-medium surface tension between polar and apolar cells was sufficient to drive internalisation (9).

Further models have examined the relative roles of position and polarisation during trophectoderm differentiation. For example, Krupinski et al. modelled this using a simplified gene network, identifying that a position-based approach for trophectoderm differentiation was more robust than a polarisation approach (17). Nissen et al. also probed the importance of polarisation for trophectoderm cells using a phenomenological agent based model, which showed that polarity dependent interactions between trophectoderm were integral for proper development (24).

However, there remain many biophysical questions that modelling has the potential to tackle. These include how the cortical tension varies around single blastomeres (along with precisely how the tension must change during compaction and internalisation), the rules governing the first sets of division (and how these differ between divisions 1-3 and divisions 4-5), the extent to which the zona pellucida is initially soft and then hardens (25–27), why the morula does not normally decompact during internalisation, and how conserved the initial developmental programme is across mammals.

Here, to answer these questions, we develop a novel computational vertex model that describes development of a mature morula from a fertilised egg. Unlike traditional vertex models, our model allows cells to separate and for the first time includes contributions from a range of realistic biophysical forces along with cell division, a zona pellucida and system noise. After first explaining the model in detail, we probe the role of cortical tension during pre-compaction. We then examine the orientation of the first two division axes and the impact of the ZP on initial cell alignment, before determining how adhesion and cortical tension are altered during compaction and internalisation. Next, we examine the role of noise in the system, followed by investigating the relative importance of position versus polarisation in deciding the fate of outer cells and probing the different division regimes during the fourth and fifth cleavages. Finally, we compare human and mouse, aiming to understand the similarities and differences of their developmental programmes.

## 2. Model development

As we recently reviewed, there are several possible modelling frameworks that could be used to describe the very earliest stages of mammalian development, from the single fertilised egg up to the mature morula (5). There are advantages and disadvantages to each, and here we choose to employ a type of 2D vertex modelling, which allows realistic biophysical forces to be implemented on the edge of each cell. Although the morula is three-dimensional, a 2D model is sufficient to investigate the most important questions (such as the cellular dynamics and the relative role of different forces) and has the advantages of being simpler to develop, quicker to simulate and easier to compare to 2D imaging data.

Vertex models describe each cell as a polygon consisting of a number of connected nodes (the vertices). Classical vertex models typically share vertices between neighbouring cells, a consequence of such models being used to describe tightly-packed epithelial tissues. However, here, in order to allow gaps to form between cells, we instead represent each cell by its own set of vertices, with vertices on neighbouring cells able to be linked via an adhesion force. Such a model has not previously been used to describe this stage of embryogenesis, although has occasionally appeared in other contexts, such as to investigate jamming in foams and emulsions or cell sorting (28, 29).

On each vertex, three or four forces are imposed corresponding to cortical tension, cell-volume conservation, membrane curvature and adhesion to neighbouring cells (see Fig. 1). The adhesion force only operates if a cell is sufficiently close to another cell. By also including a drag force on each vertex proportional to the velocity and neglecting inertial effects (as is typical in low Reynold number environments; here Re < 10^−4^), we are able to derive the velocity on each vertex (30). See the Supporting Information for full details.

**Fig. 1.**
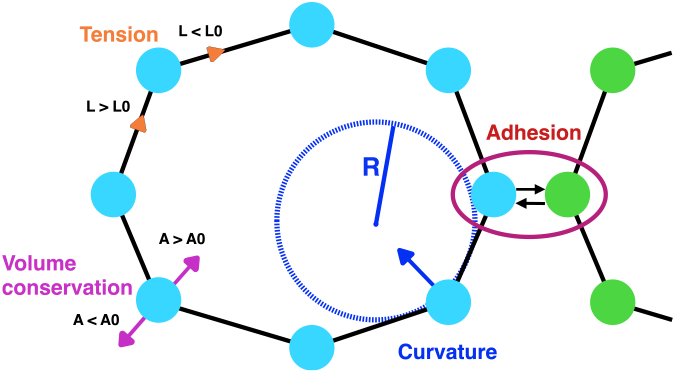
Schematic of the forces implemented on each vertex. The tension force acts directly along edges, either towards vertices if the cell circumference *L* is greater than its unstressed length *L*_0_ or away from vertices if *L < L*_0_. Similarly, the volume force acts normal to the cell surface, either inwards or outwards depending on whether the cell area *A* is greater or less than its resting area *A*_0_. Curvature acts to flatten the membrane (assuming no spontaneous curvature) with a magnitude related to the local radius of curvature *R* at each vertex. Finally, adhesion acts between two vertices on neighbouring cells that are sufficiently close to each other.

We now explain the four forces in turn. First, we consider the tension force, *F*_tension_ (23). We assume this is a global force related to conservation of the area of the cell boundary (made up of the cell membrane and underlying actin cortex). Although there is a degree of spare membrane within, for example, membrane ruffles, the cortex gives the cell boundary a degree of rigidity that resists changes to its area (31). The tension force acts in the direction of the edges between neighbouring vertices (see Fig. 1). In our model, we assume that 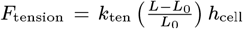, where *k*_ten_ is the tension force coefficient, *L* is the total circumference of the cell, *L*_0_ is the resting circumference of the cell and *h*_cell_ is the cell height.

Although we implement a global tension force, we allow the tension force coefficient, *k*_ten_, to be different along adhered and unadhered regions of the cell boundary. This is a distinct effect to the adhesion force discussed below and captures the different surface tensions that have been measured around the cell, likely due to differing actin structures and thicknesses beneath the adhered and unadhered regions of the cell (10). Although different parts of the cell boundary may well also have their own resting circumferences, *L*_0_, we choose for simplicity to consider a single global *L*_0_ for each cell and to allow only *k*_ten_ to change. In particular, we write *k*_ten_ as *γ* along adhered regions and as Γ along unadhered regions (see Fig. 2A).

**Fig. 2.**
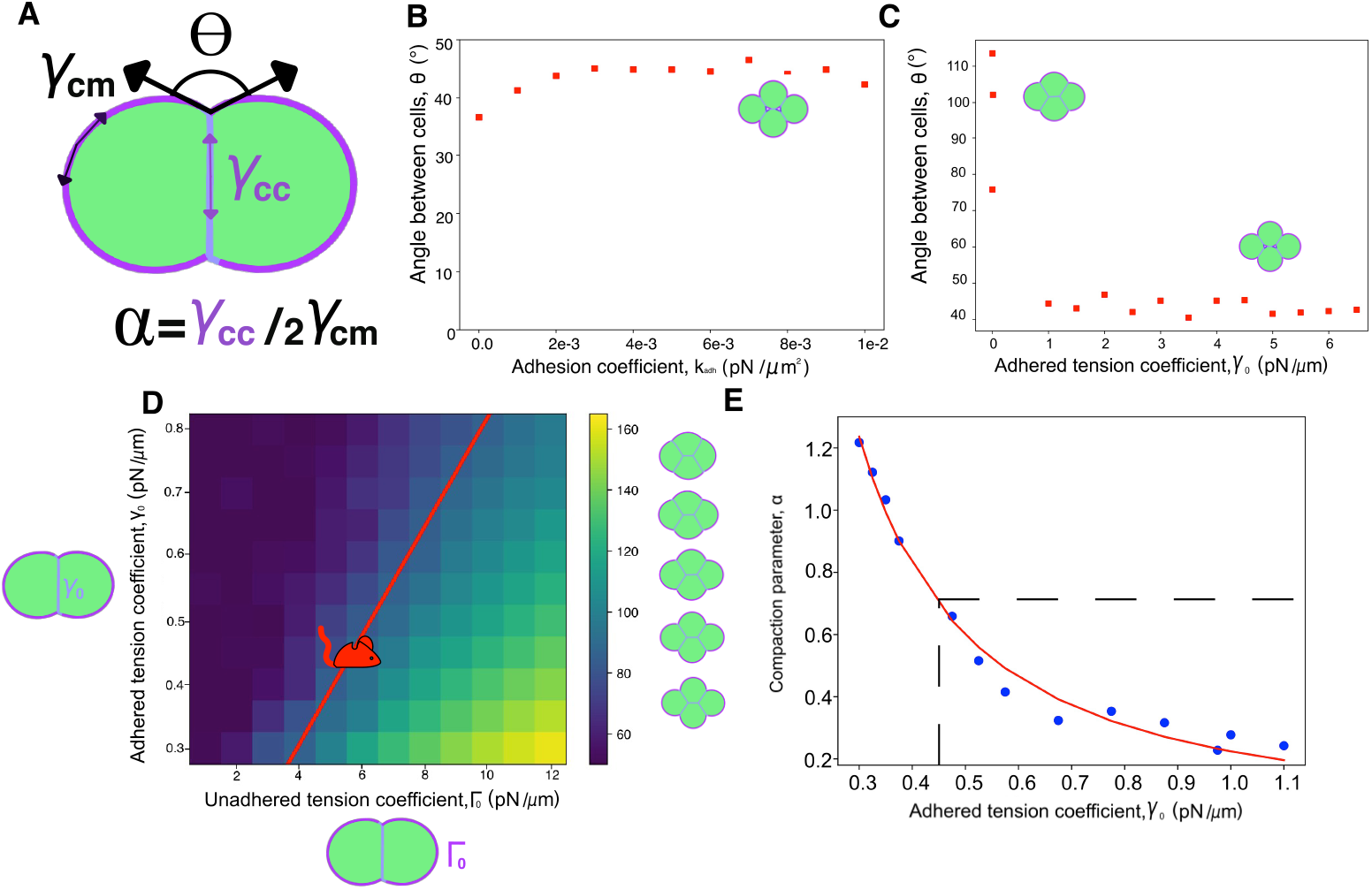
Pre-compaction results. (A) The definition of the angle between cells, *θ*, and the compaction parameter, *α*, in terms of the surface tensions at the cell-cell interface, *γ*_cc_, and cell-medium interface, *γ*_cm_. (B) The average angle between cells *θ* against the adhesion coefficient *k*_adh_ with Γ_0_ = *γ*_0_ = 0.45pN/*μ*m. Small sketches of cells in this and other panels show a typical configuration at that particular point. (C) The average angle *θ* against *γ*_0_ with Γ_0_ = *γ*_0_ and *k*_adh_ = 0.01*pNμm*^−2^. (D) Heat map of *θ* against Γ_0_ and *γ*_0_ with *k*_adh_ = 0.01*pNμm*^−2^ showing that the highest angles are associated with large values of Γ_0_ and small values of *γ*_0_. The red line shows parameter values corresponding to the experimentally-measured value of *θ* = 85^°^. The mouse symbol denotes the particular values for Γ_0_ and *γ*_0_ that we infer. (E) The surface tension ratio *α* against *γ*_0_ along the red line in (D). The red line shows the best fit to 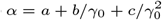. The dotted line shows the measured experimental value of *α* = 0.71 and the corresponding value of *γ*_0_ = 0.45pN/*μ*m (paired with Γ_0_ = 6pN/*μ*m from (D)).

The volume force is implemented in a similar way to the tension force, with a resting area *A*_0_ (in two dimensions the cell volume becomes an area) for each cell corresponding to an unstressed state. For a cell of area *A* this leads to 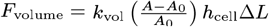, where *k*_vol_ is the compressibility constant (32) and Δ*L* is the lattice spacing. Unlike the tension force, the volume force acts normal to the cell surface. The initial cell divisions involve cleavage, where the cell does not immediately grow after division. We model this by halving *A*_0_ after each division.

For the curvature force, we first note that the cell boundary has a preferred curvature called the spontaneous membrane curvature, which for simplicity we assume to be zero. Then any deviation from a flat membrane leads to a curvature force, *F*_curvature_. Motivated by the famous Helfrich formula for the membrane curvature energy, we model the curvature force as *F*_curvature_ = *k*_curv_*h*_cell_Δ*L/R*, where *k*_curv_ is the curvature constant (related to the membrane bending modulus) and *R* is the local radius of curvature (33).

Finally, adhesion is implemented on any vertex that is within a certain capture radius, *R*_capture_, of another vertex. Such adhesion is due to proteins such as Cdh1 (34–36). For vertices within *R*_capture_ of each other, the adhesion force is implemented as a constant attractive force, *F*_adhesion_ = *k*_adh_*h*_cell_Δ*L*, acting along the line joining the two vertices, where *k*_adh_ is the adhesion coefficient.

There are a number of other components to our model that we briefly cover now. Full details are in the Supporting Information. First, cells can divide via cleavage. This occurs randomly, but such that cells divide on average between 12 and 20 hours after the last division (depending on the developmental stage) (37) and with a standard deviation of between 1-2 hours (38). There are various rules by which the division axis can be chosen, including via a random axis or by dividing along the cell’s longest axis, and we consider and compare a number of these options below. Second, we include a zona pellucida (ZP) by imposing a hard ellipsoidal shell through which vertices cannot move. Vertices are moved back to their previous position if they try to cross this shell. Third, Gaussian noise with a magnitude proportional to the square root of the time step (as per the Euler-Maruyama method) is included when required. For an example of a complete simulation output, see Supporting Movie S2.

## 3. Results

### A. Pre-compaction

We first study the very earliest stages of embryogenesis, during the initial rounds of cell division, when the total number of cells is at most eight. This occurs before the compaction stage and allows us to determine the basic parameters of our model, and in particular to study the relative sizes of the tension and adhesion forces. We initially focus on mouse and later extend our results to human.

During this pre-compaction stage, we label the tension coefficients with a subscript 0, so that *γ*_0_ is the coefficient along adhered regions and Γ_0_ the coefficient along unadhered regions. Later we will use a subscript 1 to denote these coefficients during compaction and a subscript 2 during internalisation.

There are two sets of pre-compaction experimental data that we use in this section, both measured by Maître et al.: the measured average angle between cells *θ* and the compaction parameter *α* = *γ*_cc_/2*γ*_cm_ where *γ*_cc_ is the surface tension at cell-cell interfaces and *γ*_cm_ the surface tension at cell-medium interfaces (see Fig. 2A). See the SI for how we calculate *θ* and *α* in our model. For mouse, the measured values are *θ* = 85^°^± 19^°^ and α = 0.71 ± 0.2 (10). We will consider the human case later.

At this stage we run our 2D simulations with four cells, as we estimate this corresponds most closely to the eight-cell stage of the real three-dimensional embryo (see the SI for details). A certain minimum tension coefficient is needed to prevent cells from ruffling, which we find to be around 0.3pN/*μ*m for both *γ*_0_ and Γ_0_.

First, we explore whether it is possible that Γ_0_ has the same value as *γ*_0_. Using a typical literature value for the tension coefficient of Γ_0_ = *γ*_0_ = 0.45pN/*μ*m, we vary the adhesion constant *k*_adh_ (10, 39, 40). Although increasing *k*_adh_ leads to a slightly increased angle between cells *θ*, we find this behaviour plateaus at an angle below 50^°^, well below the observed value of 85^°^ (see Fig. 2B). This is because, although adhesion brings adhered vertices closer together, and so brings neighbouring unadhered vertices closer together, there is a limit to this effect: when the adhered vertices are effectively on top of each other, further increasing adhesion makes practically no difference to the angle between cells. This seems to agree well with the suggestion by Maître et al. that the role of adhesion is simply to couple adjacent cell surfaces together (41).

We now examine whether fixing the adhesion constant to a typical value of *k*_adh_ of 0.01pN/*μ*m^2^ (41, 42) and instead altering Γ_0_ = *γ*_0_ can give an angle of 85^°^ (see Fig. 2C). This is indeed possible for small values of the tension coefficient, but when such a tension coefficient is used the ratio of the surface tensions between adhered and unadhered parts of the cell *α* ≈ 400 does not match the measured value of 0.71. Our conclusion is thus that there needs to be a difference in tension coefficient around the cell, agreeing well with the measured results of Maître et al. (10).

We now investigate the effect of such a difference in tension coefficients by plotting the average angle between cells as a function of Γ_0_ and *γ*_0_. As explained above, we only consider *γ*_0_ and Γ_0_ *>* 0.3pN/*μ*m to avoid ruffling. We obtain angles in the range 55^°^ to 160^°^, with the largest angles associated with small values of *γ*_0_ or large values of Γ_0_ (see Fig. 2D).

To understand this dependence on the tension coefficients, first consider the effect of decreasing *γ*_0_, the tension coefficient along adhered parts of the membrane. This reduces the inward tension in the adhered region, leaving an overall outward force that slightly expands the cell in this region. In turn this slightly flattens the membrane at the edge of the adhered region, reducing the resultant inward tension force and so slightly increasing the overall outward force in this region. In response, the part of the membrane just outside the adhered region moves outwards, increasing the adhered region size and so increasing the *θ* angle.

Next, consider instead the effect of increasing Γ_0_, the tension coefficient along unadhered regions. The reason this leads to an increased value of *θ* is more subtle. An increased Γ_0_ leads to an increased inward tension force along the unadhered part of the cell, which leads to slightly smaller cell volume. This increases the outward volume force all around the cell, which results in an overall outward force in the region adjacent to the adhered part of the cell. As with the case of decreasing *γ*_0_, this causes the membrane to move outwards at the edge of the adhered region, increasing the size of the adhered region and so increasing *θ*.

Since the observed pre-compaction angle is *θ* = 85^°^, we next found all the values of Γ_0_ and *γ*_0_ corresponding to this angle (red line in Fig. 2D). To determine which particular one of these values to use, we calculated the value of the compaction parameter *α* along this line (Fig. 2E), which shows that, to match with the measured value of *α* = 0.71 (10), we need Γ_0_ = 6pN/*μ*m and *γ*_0_ = 0.45pN/*μ*m.

Finally, if two cells are isolated from an eight-cell cluster, it has been found that the angle between cells decreases by around 10^°^ (42). We can also test this in our simulations and reassuringly find a similar decrease from around 85^°^ to 75^°^. The slightly smaller decrease may be related to our simulation being two-dimensional: extracting two from eight cells is studied in our simulation by extracting two cells from only four cells, perhaps leading to a less dramatic decrease in angle.

### B. The first divisions

Although the zona pellucida (ZP) is often measured to be spherical, in may cases it is slightly elliptical in shape (43). After the first division, when the single fertilised egg becomes two cells, it is observed that the two cells typically align along the long axis of the ZP, as shown in right-hand sketch in Fig. 3A (44). It is an open question as to whether this is due to the cells rotating in a rigid ZP (45) or due to a soft, malleable ZP that gradually reshapes around the divided cells (46). We can test this with our model by predicting how long cells would take to reorientate themselves in a hard ZP. To do this, we consider a hard, elliptical ZP with the ratio between the short-to-long-axis lengths being *χ*. We start our simulation with a single cell, which then divides in such a way that the cells are perfectly misaligned with the ZP, *i*.*e*. such that the angle between the line joining the centre of the cells and the short axis of the ZP is *ϕ* = 0^°^ (see Fig. 3A). We then run the simulation for time *T* and measure the new angle *ϕ* at the end. For realistic values of *χ* and T, there is at most modest rotation of the cells, with the final value of *ϕ* never being even close to 90^°^. We can achieve full rotation only in biologically-unrealistic situations: either with cells that are much firmer (*i*.*e*. higher cortical tension) than expected at the pre-morula stage, with extremely high levels of noise, with a ZP that is much more elongated than is observed, or after many days (see Fig. 3B). Thus our model predicts it is unlikely that cells rotate within a rigid ZP: either the fertilised egg always divides along the ZP’s long axis or the ZP is relatively soft at this stage and is able to remould itself around the two cells.

**Fig. 3.**
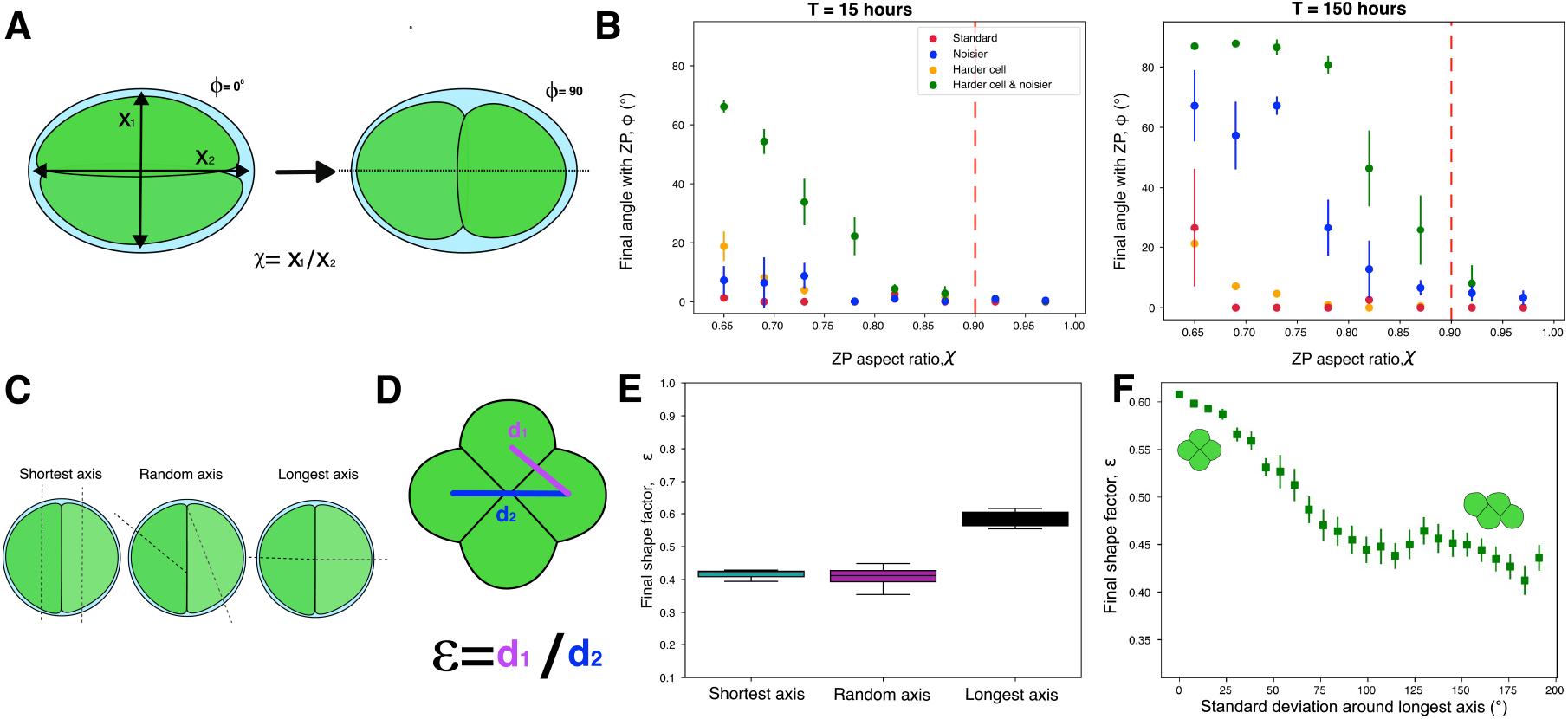
First division results. (A) Definition of *χ*, the ratio of the short-to-long-axis lengths for the ZP, and *ϕ*, the angle between the line joining the cell centres and the ZP’s short axis. (B) The final *ϕ* angle against *χ* after 15 hours (left) and 150 hours (right). Four cases are considered: standard parameters, hard cells, increased noise level, and both hard cells and noisier. The red dashed line shows the experimentally-measured value of *χ*. (C) The division scenarios that we consider: division along the shortest axis, division at a random angle and division along the longest axis. (D) Definition of the shape factor ε, the ratio of the shortest-to-longest cell-to-cell distances. (E) The final shape factor after two rounds of division, showing that only division along the longest axis can achieve a high value of ε. (F) How the average final shape factor depends on the variation of the division axis around the longest axis, showing high values of only up to a standard deviation of around 25^°^.

After the two-cell stage, both cells divide again, normally into a tetrahedron shape (47), although more planar configurations are seen in around 20% of cases (48, 49). As with the first division, the rules that govern this next set of divisions are not yet fully understood. Some have suggested that cells tend to divide always along their long axis (*i*.*e*. Hertwig’s rule) (50), whereas others argue that this is not always strictly the case (51). Although our two-dimensional model can not fully capture the three-dimensional tetrahedral configuration, we can still investigate this question by considering four possible division rules: division along the longest axis, division along the shortest axis, a random-division plane, and division around the longest axis with a Gaussian distribution (see Fig. 3C). To quantify the results, we define a shape factor, ε, for the four-cell stage as the ratio of the shortest-to-longest cell-to-cell distance (see Fig. 3D). For a perfect tetrahedron in 3D, this would give *ε* = 1. However, in our 2D model the maximum possible value is 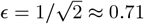. Each simulation starts from two cells, allows each cell to divide once and then measures *ε* just before the next division. Our results (see Fig. 3E) show that division along the longest axis results in the largest *ε* values, close to the theoretical maximum. Division instead along the shortest axis or a random division axis only leads to *ε* around 0.4. If we run our simulation without a ZP, we find a small drop in *E* in the case of longest-axis division, matching the model from Giommana and Campàs (18). Finally, we quantify the tolerance around long-axis division in order to still maintain a high final shape factor. To do this, we choose the division angle from a Gaussian distribution with mean along the longest axis and a given standard deviation. A high final value of *E* is still obtained for standard deviations up to about 25^°^, suggesting a reasonable variation in longest-axis division can easily be tolerated (see Fig. 3F).

### C. Compaction

During compaction, which occurs in mice at the eight-cell stage, the cell-to-cell contact area is increased, which leads to an associated increase in the angle between cells. In mouse, Maître et al. found this increase to be from 85^°^ to 146^°^, along with a decrease in *α* from 0.71 to 0.25 (10).

One of the first mathematical models of this process proposed that compaction is caused by differing cell surface tensions at cell-to-cell contacts compared to exposed cell surfaces (19). Building on this idea, there are two main hypotheses that might account for this change in cell surface tension and so be the driving force of compaction: differential adhesion (*i*.*e*. increased adhesion between cells) and differential interfacial tension (*i*.*e*. decreased tension along cell-to-cell contact regions or increased tension along non-adhered regions). It is now generally thought to be a combination of the two mechanisms (52, 53).

Maître et al. recently found that increases in surface tension in mouse embryos at the cell medium interface as well as decreasing surface tension at the cell-to-cell boundary were mostly due to actin-myosin contractility (10), with seemingly little contribution from adhesion effects. In addition, it has been suggested that E-cadherin-dependent filopodia also play a role at the cell-medium interface, although this has been disputed (10, 34).

Previous models of compaction were energy-based and did not differentiate between cortical tension and adhesion contributions towards surface tension. Here, with our force-based approach, we probed these effects separately (9, 19, 54). First, with the values of Γ_0_ = 6pN/*μ*m and *γ*_0_ = 0.45pN/*μ*m found for pre-compaction, we investigated whether compaction can be achieved simply by increasing the adhesion coefficient. However, this showed very little change in angle (see Fig. 4A), agreeing with both the experimental and computational results in Maître et al. (10). As in pre-compaction, this is likely because increased adhesion simply moves already-close vertices slightly closer together. Adding noise to the vertex position does not change this conclusion: noise encourages more vertices to adhere, but also causes adhered vertices to separate, making practically no overall difference.

**Fig. 4.**
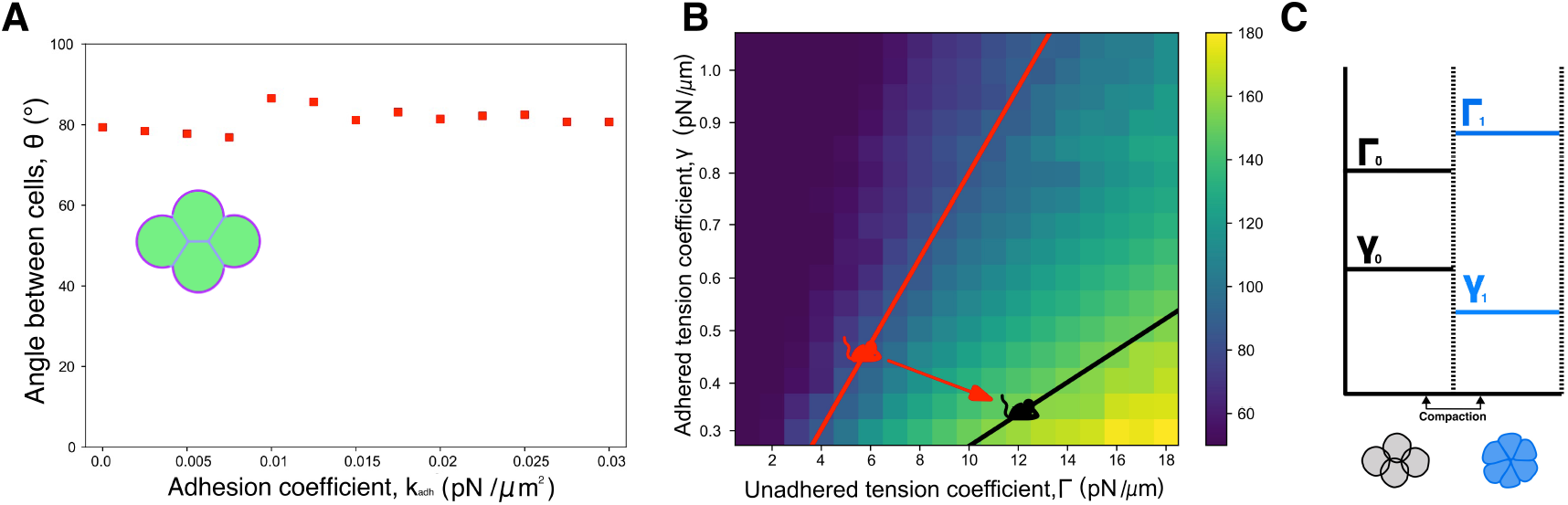
Compaction results. (A) The average angle between cells *θ* versus the adhesion coefficient *k*_adh_ with Γ_0_ = 6pN/*μ*m and *γ*_0_ = 0.45pN/*μ*m as found in the pre-compaction section. (B) Heat map of *θ* against Γ and *γ* with *k*_adh_ = 0.01*pNμm*^−2^. The red line shows the values of Γ_0_ and *γ*_0_ giving a pre-compaction angle of *θ* = 85^°^. The black line shows the values of Γ_1_ and *γ*_1_ giving a post-compaction angle of *θ* = 146^°^. The mouse symbols show the actual values of Γ_0_ and *γ*_0_ (red mouse) and Γ_1_ and *γ*_1_ (black mouse) that we infer from the measured compaction parameters *α*. (C) Schematic (not to scale) showing how the two tension coefficients change in opposite directions during compaction.

Next, we considered whether compaction can be explained by a change in one of the tension coefficients. We do this by changing the coefficient at the start of compaction from *γ*_0_ to *γ*_1_ along adhered regions or from Γ_0_ to Γ_1_ along unadhered regions. We found that increasing just the Γ value to Γ_1_ = 16pN/*μ*m^−1^ (with no change in *γ*) gives the observed angle of 146^°^ (Fig. 4B), whereas just decreasing the *γ* value (with Γ fixed) cannot achieve this unless an unphysically small *γ*_1_ value is used.

However, it has been suggested that in fact both Γ and *γ* are changed during compaction (see Fig. 4C) (10). By also using the measured decrease in *α* (as in the pre-compaction section), we were able to confirm this and predict that during compaction Γ increases to Γ_1_ = 12pN/*μ*m and *γ* decreases to *γ*_1_ = 0.35pN/*μ*m (see Fig. 4B).

Finally, as with the pre-compaction case, we used these values of Γ_1_ and *γ*_1_ to investigate the change in angle if two cells are extracted from a compacted morula, finding a decrease in *θ* from 146^°^ to 135^°^. Reassuringly, this is again similar to the measured change of around 15^°^ found in de Plater et al. (42).

### D. Internalisation

After compaction, the morula undergoes internalisation, where a subset of cells move to the centre so they are completely surrounded by other cells, with no part of their surface on the outside. To do this, cells are split into either polar cells (which remain on the outside) or apolar cells (which internalise). This is accomplished by Pard-6 localising to the exposed outer surface in polar cells, leading to F-actin being redistributed (55, 56). In particular, in polar cells, Pard-6 negatively regulates the actomyosin network resulting in an apical domain that is depleted of actin and surrounded by an apical ring (9, 57). It is believed that this difference in the actin network between polar and apolar cells leads to a difference in surface tensions and so to internalisation (9, 35).

To describe this situation in our model, we split cells into polar cells (with tension coefficients Γ_2,polar_ and *γ*_2,polar_) and apolar cells (with tension coefficients Γ_2,apolar_ and *γ*_2,apolar_). In particular, we start from a symmetric arrangement of six cells with one apolar cell (which will try to internalise) and five polar cells (see Fig. 5A). This is the fewest number of cells in our simulation where complete internalisation can be observed and matches well with the observation that the first cell internalisation most often happens before the 12-cell embryo (35). To quantify internalisation of the apolar cell, we define its degree of internalisation, Ψ, as the ratio of the length of its boundary adhered to other cells to its total boundary length. A value of Ψ = 1 corresponds to complete internalisation, while Ψ = 0 would represent a cell completely detached from any other cell. Our initial arrangement of six cells corresponds to Ψ = 0.71.

**Fig. 5.**
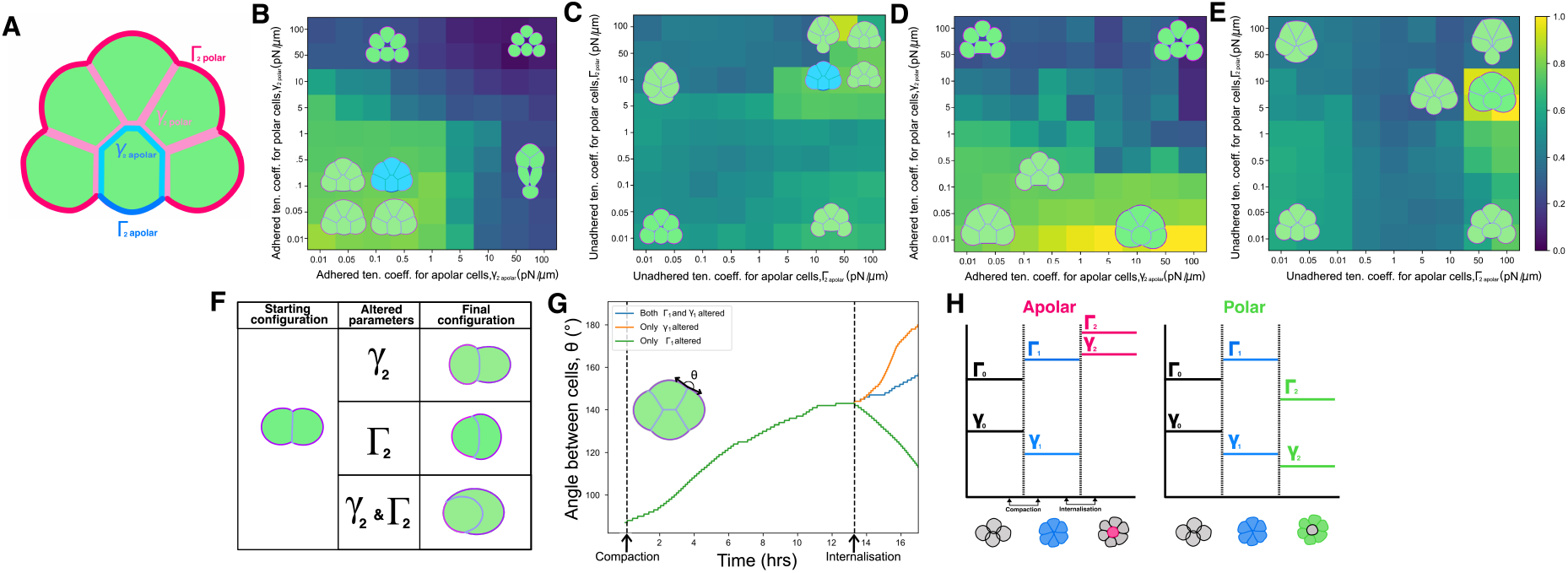
Internalisation results. (A) Sketch of the four tension coefficients at the internalisation stage: two for polar cells (Γ_2,polar_ and *γ*_2,polar_) and two for apolar cells (Γ_2,apolar_ and *γ*_2,apolar_). (B) Degree of internalisation Ψ against the two adhered tension coefficients (*γ*_2,apolar_ and *γ*_2,polar_) with no change in the unadhered tension coefficients (Γ_2,apolar_ = Γ_2,polar_ = Γ_1_). The blue sketch shows the initial configuration. (C) Degree of internalisation Ψ against the two unadhered tension coefficients (Γ_2,apolar_ and Γ_2,polar_) with no change in the adhered tension coefficients (*γ*_2,apolar_ = *γ*_2,polar_ = *γ*_1_). (D) As in (B) but with the unadhered tension coefficients changed to Γ_2,apolar_ = 100pN/*μ*m and Γ_2,polar_ = 4pN/*μ*m. (E) As in (C) but with the adhered tension coefficients changed to *γ*_2,apolar_ = 30pN/*μ*m and *γ*_2,polar_ = 0.01pN/*μ*m. (F) Results for a two-cell doublet, showing envelopment only when both Γ and *γ* are changed. (G) How the angle between cells *θ* changes over time from pre-compaction through compaction and then internalisation. Compared are three internalisation scenarios: only Γ values are changed, only *γ* values are changed, and both Γ and *γ* values are changed). The first dashed vertical line shows the start of compaction when Γ_0_ → Γ_1_ and *γ*_0_ → *γ*_1_. The second dashed vertical line shows the start of internalisation when Γ_1_ → Γ_2_ and *γ*_1_ → *γ*_2_. (H) Schematic (not to scale) showing how the tension coefficients change during compaction and internalisation for both apolar and polar cells.

We first investigate whether a difference purely in *γ* (*i*.*e. γ*_1_ splits into *γ*_2,polar_ and *γ*_2,apolar_; Fig. 5B) or in Γ (*i*.*e*. Γ_1_ splits into Γ_2,polar_ and Γ_2,apolar_; Fig. 5C) can instigate internalisation. Although this leads to the apolar cell partially internalising for some parameter values, the lack of spreading of the neighbouring cells over its exposed surface hinders full internalisation. In order to have full internalisation (Ψ = 1), we need both *γ*_1_ and Γ_1_ to change from their values during compaction (Figs. 5D and E), with the highest level of internalisation associated with *γ*_2,polar_ = 0.01pN/*μ*m, *γ*_2,apolar_ = 30pN/*μ*m, Γ_2,polar_ = 4pN/*μ*m and Γ_2,apolar_ = 100pN/*μ*m (see sketch in Fig. 5H).

It is also possible to investigate internalisation (perhaps better called envelopment) of just two cells, one polar and one apolar. This was studied by Anani et al. by isolating a single polar cell from an eight-cell blastomere and allowing it to divide. Cases of asymmetric division created such a polar-apolar doublet, which showed the apolar cell being enveloped by the polar cell (58). In our simulation we find that only splitting *γ*_1_ or only splitting Γ_1_ does not lead to doublet engulfment. Rather, we need to split both *γ*_1_ and Γ_1_ to the above four values to observe envelopment (see Fig. 5F).

Finally, we examine the potential for decompaction. Small levels of decompaction and subsequent recompaction have been observed in both mouse and human, but this is not the norm (59, 60). Since Γ_0_ is increased to cause compaction and then decreased for internalisation, there is the possibility that the cells will then start to decompact. We study this in our model by examining four polar cells and plotting the outside angle as a function of time for different changes to *γ*_1_ and Γ_1_ (Fig. 5G). If, at the point the cells start to internalise, only Γ_1_ is decreased to Γ_2,polar_ (with no change in *γ*_1_) then we do indeed observe decompaction. However, if instead (or as well) *γ*_1_ is decreased to *γ*_2,polar_, then no decompaction occurs. In fact, in these cases, our model predicts that cells start to compact even more. As far as we know, no attempt has yet been made to measure this and it would be interesting to determine if the angle *θ* for polar cells does indeed continue to increase during internalisation.

### E. The role of noise

Up until now there has been no noise in our simulations except for the stochasticity in cell division timing. However, it has been suggested that noise may play an important role during early development, especially during internalisation and cell sorting (17). There are a number of ways that we could include a noise component to our model, but here we add a noise force to each vertex that points in the normal direction and has magnitude 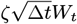, where *ζ* is the noise coefficient, Δ*t* the time step, and *W*_*t*_ is drawn from a Gaussian distribution with mean 0 and standard deviation 0.2 (61).

We first examine the effect of noise on compaction, finding that the final angle between cells decreases with increased noise (Fig. 6A). This is likely because noise is disrupting cell-to-cell adhesion, leading to a smaller adhered region between cells. Noise also has a significant effect on the compaction time, with large levels of noise taking over 25% longer to reach full compaction (Fig. 6A).

**Fig. 6.**
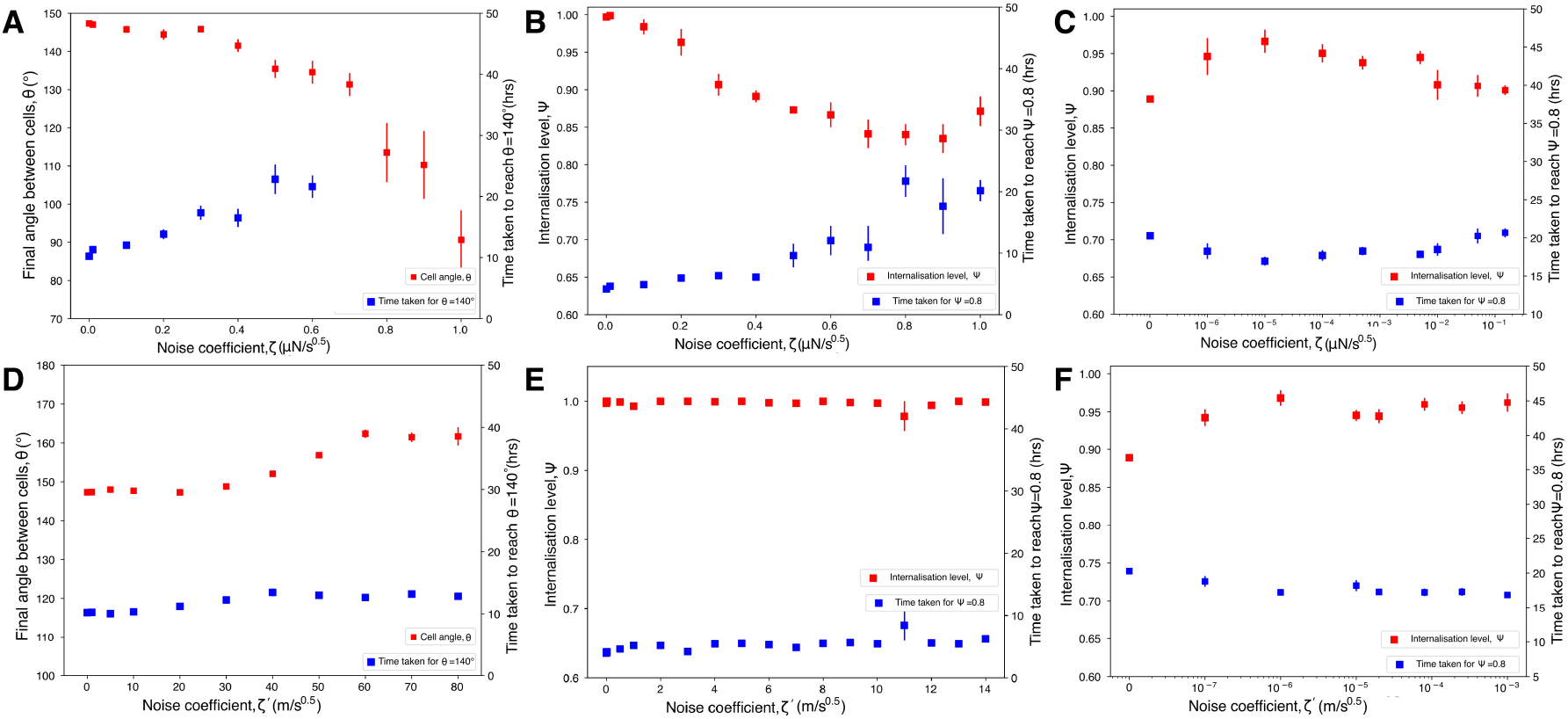
The effect of noise. (A) Final angle between cells *θ* (red) and time taken to reach *θ* = 140^°^ (blue) against the noise coefficient *ζ*, showing that increased noise leads to lower angles between cells and longer compaction times. (B) Internalisation level Ψ (red) and time taken to the reach Ψ = 0.8 (blue) against the noise coefficient *ζ*, showing that higher noise leads to lower internalisation and longer internalisation times. (C) As in (B) but starting from tension coefficients that are 90% of their optimal values. (D-F) As for (A-C) but with active noise (and so with the modified noise coefficient *ζ*^*1*^).

The situation is similar for internalisation. Noise hinders the internalisation process, with increased noise leading to lower degrees of internalisation, Ψ (Fig. 6B). Cells also internalise slower with increasing noise, as the apolar cells are impeded as they try to move towards the centre of the morula. These results are for the optimal tension coefficients found above. We can also start our simulation from non-optimal parameter values to investigate whether noise is able to internalise a cell that would not otherwise internalise. To do this, we change each of the four tension coefficients by 90% in the direction of their compaction value. This shows that small noise levels are able to help internalisation for non-optimal tension coefficients, but that too much noise starts to have a detrimental effect (Fig. 6C).

These results correspond to “passive” noise, where the size of the noise *ζ* is independent of any local properties. However, Yanagida et al. measured different cell surface fluctuations between epiblast and primitive endoderm cells, suggesting that noise may differ for distinct cell types and at different points around the cell (62). While not a direct comparison of the blebbing discussed by Yanagida et al., we choose to implement this “active” noise by letting the noise coefficient be directly proportional to the local cortical tension *k*_ten_ so that *ζ* = *ζ*^′^*k*_ten_ for a new noise coefficient *ζ*^′^. In addition, we now let the noise force point in the same direction as the tension force rather than in the normal direction.

For compaction, active noise exhibits very different behaviour to the passive noise case. Now increasing levels of noise lead to a larger compaction angle (Fig. 6D). This is likely because unadhered parts of the membrane now have increased noise over the adhered parts. This means that unadhered vertices are more likely to find a vertex on a neighbouring cell to adhere to, and are then less likely to unadhere once they have adhered. One consequence of this is that active noise requires smaller changes in Γ and *γ* to obtain the same final compaction angle, so that a less dramatic change to the actin cytoskeleton can cause the same level of compaction.

For internalisation with the optimal tension coefficients, active noise makes little difference to either Ψ or the time taken to reach it (Fig. 6E). However, if we again change the optimal parameters by 90% so that internalisation is no longer observed in the absence of noise, then a small level of active noise is again able to recover internalisation, potentially providing a degree of stability to the system (Fig. 6F). Further, larger levels of noise now maintain a high Ψ value, even in situations where passive noise would start to become detrimental.

### F. The fourth and fifth divisions

After three division cycles the morula contains eight cells. At this stage, the inner cell mass (ICM) and trophectoderm (TE) start to form over the next two rounds of division. The rules controlling these divisions are still a topic of debate, in particular the question of how TE fate is decided. There are two main competing ideas—that cells become TE based on their position or based on their polarisation (*i*.*e*. whether they contain an apical domain)— with the latter the more widely accepted (12, 17, 63–66). Inheritance of the apical domain after the eight-cell stage has been shown to induce TE specification, with the proportion of the apical domain inherited being linked to the level of TE gene expression (13).

To test the relative importance of position-versus-polarisation in TE fate, we ran our simulation starting from five polar cells arranged in a circle, each with an apical domain on the outside (see Fig. 7A). We then allowed one of the cells to divide along a randomly-chosen axis and tracked the internalisation level, Ψ, of one of its daughters (chosen at random) over time.

**Fig. 7.**
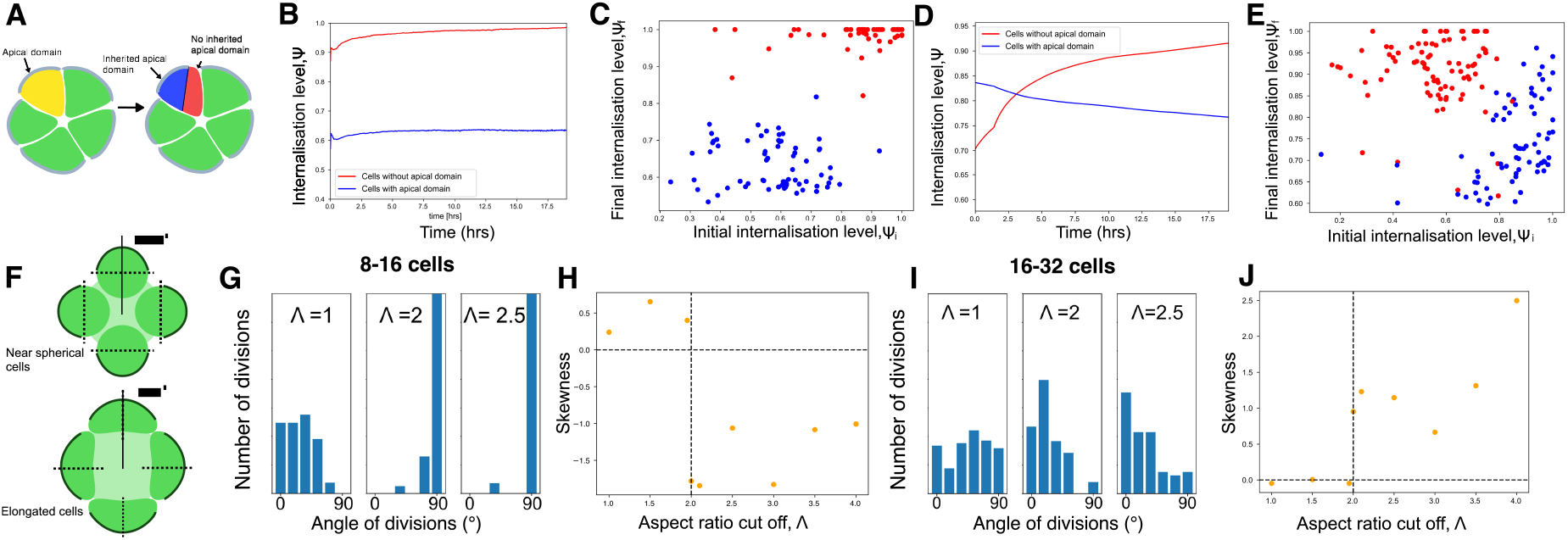
Results for later divisions. (A) Sketch of how, after a polar cell divides, the apical domain (shown in grey) is not necessarily inherited by both daughter cells. Here, after division of the yellow cell, only the blue daughter cell inherits the apical domain. (B) The average internalisation level Ψ as a function of time for the case that, upon division, the most external daughter cell inherits the apical domain. Both daughter cells are tracked, with the inside cell (without the apical domain; shown in red) showing complete internalisation, whereas the outside cell (with the apical domain; shown in blue) remains uninternalised. (C) Scatter plot of the initial internalisation level Ψ_*i*_ against the final level Ψ_*f*_ for the same situation as in (C). The daughter cells with an apical domain (blue) start and remain on the outside, whereas the daughter cells without an apical domain (red) almost all end up completely internalised. (D) Same as (B) but for the case that the inside daughter cell is the one that inherits the apical domain, demonstrating that the two cells switch positions, with the daughter cell without the apical domain becoming internalised. (E) Same as (C) but for the case shown in (D). (F) Sketch of the possible division axes, with near-spherical cells dividing so that one daughter inherits the whole apical domain (asymmetric division) and more elongated cells dividing along their longest axis (symmetric division). The division angle Ω measures the angle between the division plane and the line joining the cell centre to the centre of the morula. (G) Distributions of the division angle Ω at the 8-16-cell stage for three aspect ratio cut-offs Λ. (H) The skewness of the distributions in (G) against the aspect ratio cut-off Λ, showing a sharp switch to negative skew around Λ = 2.0. (I) As in (G) but for the 16-32-cell stage. (J) As in (H) but for the 16-32-cell stage, which now shows a sharp increase to positive skew at the same value of Λ = 2.0.

We first consider the case that the daughter cell on the outside inherits the apical domain, with the internal cell becoming apolar (as in Fig. 7A). As expected, this shows that the average level of internalisation remains roughly constant over time (Fig. 7B). Cells that start on the outside remain on the outside, and cells that start towards the middle become completely internalised (Fig. 7C).

Then we tried switching which daughter cell receives the apical domain so that, unlike in Fig. 7A, it is the most-internal cell that inherits the domain. Although unlikely to be biologically plausible, this allows us to investigate an extreme case to determine the relative roles of position and polarisation during internalisation. Interestingly, our model shows that the daughter cells completely switch positions after only a couple of hours (Fig. 7D), with the initially-outside cell without the apical domain being the one that becomes internalised (Fig. 7E). This strongly suggests that cell polarisation (*i*.*e*. the presence or absence of an apical domain) is the dominant factor determining which cells internalise, with cell position of little importance for the final cell fate.

Next we examine how the division axis is chosen at the 8-to-32-cell stage. It has been suggested that during these two sets of divisions the division axis is chosen in one of two ways: either “symmetric division” where cells divide along their longest axis (*i*.*e*. Hertwig’s rule) or “asymmetric division” where cells divide so that the apical domain is inherited by only one daughter cell (*i*.*e*. the apical domain is asymmetrically segregated). Niwayama et al. claim that the choice of symmetric or asymmetric division is related to the cell morphology, with elongated cells dividing symmetrically and more spherical cells dividing asymmetrically (see Fig. 7F) (67).

To study this we started our simulation from four cells (representing eight cells in 3D) and allowed them to divide twice each until 16 cells appear (representing 32 cells in 3D). For each division, we define the division angle Ω as the angle between the division plane and the line joining the cell centre to the centre of the morula (see Fig. 7F). We then introduced an aspect ratio cut-off Λ such that a cell divides symmetrically if the ratio of its longest-to-shortest axis is greater than Λ and asymmetrically if the ratio is less than Λ. Running our simulation many times gives a distribution of division angles Ω for each value of the cut-off Λ.

For the 8-16-cell stage we found a marked difference in the distribution of division angles for different cut-offs. At low values of Λ (where almost all cells divide symmetrically) the division angle Ω is biased towards small angles. However, for large Λ (where almost all cells divide asymmetrically) Ω is clustered around *θ* = 90^°^ (Fig. 7G). To quantify this switch, we plotted the skewness of the division angle distribution against the cut-off, which shows an sudden sharp decrease to negative skew around Λ = 2.0 (Fig. 7H). This distribution was experimentally measured by Watanabe et al., who found a negatively-skewed distribution (12). (Note that our definition of the division angle differs to that in Watanabe et al.: Ω_here_ = 90^°^− Ω_Watanabe_.) Thus our model predicts that Λ > 2, *i*.*e*. cells whose long axis is more than twice their short axis divide symmetrically, whereas other cells divide asymmetrically.

For the 16-32-cell stage we found a dramatic difference to the 8-16-cell stage. Now for low cut-offs the division angles show a fairly uniform Ω distribution. When Λ is increased this distribution becomes more clustered around small division angles, exactly the opposite of the 8-16-cell stage (Fig. 7I).

The plot of skewness against cut-off confirms this, with a switch from zero to positive skew as Λ is increased (Fig. 7J). Interestingly, this transition is again remarkably sharp and occurs for exactly the same cut-off of Λ = 2.0. These results again agree with Watanabe et al., who measured a shift from a negative to positive skew distribution between the 8-16 and 16-32-cell stages (12).

### G. Human versus mouse

We have so far focussed on mice by exclusively fitting our parameters to mouse data. However, one advantage of our model (and modelling approaches in general) is that it is easily adapted to other organisms. We now do this by applying our model to human embryogenesis and so demonstrate that the mechanisms and processes underlying early development of mouse and human are similar, particularly with regard to the biophysics behind compaction and internalisation.

Mice and humans share many similarities during the first few stages after fertilisation, including compaction and internalisation, and it has been suggested that the mechanisms of polarisation are mostly conserved between the two species (68, 69). However, there are also notable differences. For example, the fertilised human egg is 50% larger than that for mouse. Also, unlike in mouse, trophectoderm factors in human are expressed independently of the polarity machinery (68, 70).

One of the chief other differences between mouse and human is that of timescales, with compaction and internalisation in mouse taking around 12 hours and starting at the eight-cell stage, whereas in human they take about a day and start at the 16-cell stage (68). The reason for the delay in human compaction is not fully understood, but is likely to be related to distinct biophysical properties of blastomeres in human compared to mouse (68). In our simulation, this difference in timescales can be captured either by changing the frictional coefficient, *η*, or by altering the average time between divisions.

Another way in which mouse and human differ during preimplantation development is in the precise angles between cells *θ* and the compaction parameter *α*. For mouse, the average *θ* increases from 85^°^ to 146^°^ during compaction, with *α* changing from 0.71 to 0.25 (10). This contrasts with the human results, where *θ* increases from 81^°^ to 158^°^, while *α* decreases from 1.00 to 0.25 (22). We can use these data to infer the tension coefficients for human in exactly the same way we did for mouse. This predicts that both pre-compaction tension coefficients are about 1.5 times larger in human, with *γ*_0,human_ = 0.75pN/*μ*m and Γ_0,human_ = 8.5pN/*μ*m (cf. *γ*_0,mouse_ = 0.45pN/*μ*m and Γ_0,mouse_ = 6pN/*μ*m) (see Fig. 8A). This agrees with the experimental results by Firmin et al., who measured the cell surface tension in human to be larger than that in mouse (22). As with mouse, our model shows that in human both

**Fig. 8.**
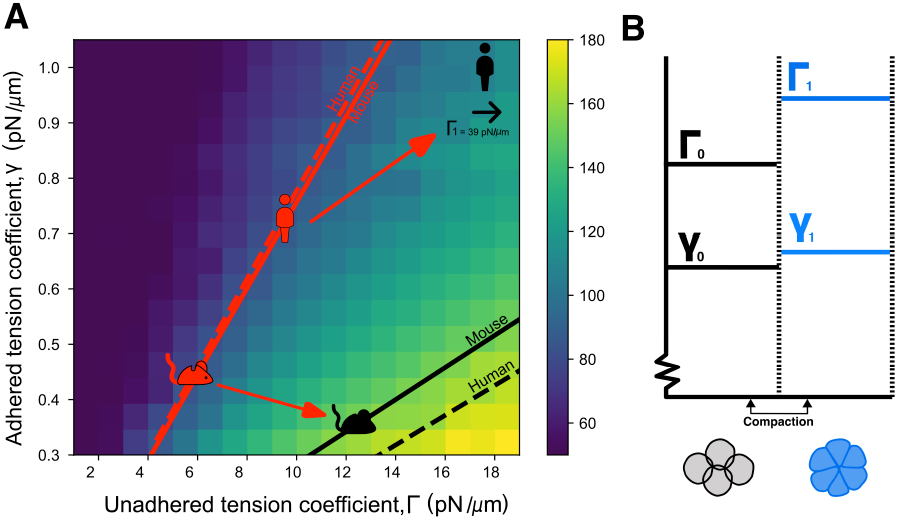
Comparison of human and mouse. (A) Heat map of the angle between cells *θ* against the adhered and unadhered tension coefficients Γ and *γ*. The red lines show points corresponding to experimentally-measured pre-compaction angles (85^°^ for mouse (solid) and 81^°^ for human (dashed)). The black lines show points corresponding to experimentally-measured post-compaction angles (146^°^ for mouse (solid) and 158^°^ for human (dashed)). The mouse and human symbols show the inferred Γ and *γ* values in each case, found by fitting to the relevant compaction parameter, *α*. (B) Schematic (not to scale) showing how the two human tension coefficients change during compaction. Unlike for mouse where Γ and *γ* change in opposite directions, our model predicts that both Γ and *γ* increase during compaction in humans.

Γ and *γ* change during compaction. However, whereas Γ increases and *γ* decreases in mouse, we predict that both tension coefficients increase in human, with *γ* increasing to *γ*_1,human_ = 1.1pN/*μ*m, whereas Γ is significantly increased to Γ_1,human_ = 39pN/*μ*m (cf. *γ*_1,mouse_ = 0.35pN/*μ*m and Γ_1,mouse_ = 12pN/*μ*m) (see Figs. 8A and B). This also agrees with Firmin et al., where they show a slight increase in cell-cell surface tension and a much larger increase in cell-medium surface tension in human (22). It is not clear why human and mouse development differs in this way, although it is tempting to speculate that it is related to an *r/K* selection effect: human embryos spend more time and energy increasing the unadhered tension constant Γ during compaction, with a corresponding larger angle between cells *θ* and a subsequent greater chance of successful internalisation.

## 4. Discussion

We developed a novel vertex model of the very earliest stages of the mammalian embryo, from the moment the egg is fertilised up to the mature morula. Unlike vertex models of epithelial sheets, vertices in our model are not shared between neighbouring cells, allowing cells to separate from each other and intercellular gaps to form. We include all known relevant biophysical forces, in particular cortical tension, membrane curvature, cytoplasm compressibility and cell-to-cell adhesion. Further, we include a zona pellucida (ZP), cell division and the effect of a noisy environment.

The first key finding of our model is that, even during the first few divisions, the cortical tension must differ around the cell. Simply including a cell-to-cell adhesive force cannot account for the experimentally-observed angles between cells. The origin of this tension difference is unlikely to be the membrane itself (*e*.*g*. by altered membrane composition), but the structure of actin cortex beneath the membrane. This agrees well with the results of Maître et al., who observed pulsed contractions of the actomyosin cortex during compaction (10), and with findings at later developmental stages, particularly internalisation and subsequent cell sorting, where a non-homogeneous actomyosin network was measured along adhered and unadhered cell regions (35, 55, 57).

Next, we examined the ZP, focusing on its role during the first cell division, when the two cells are typically found to lie along the ZP’s long axis. It is still debated whether cells at this stage are able to rotate within the ZP. What our model suggests is that, although cells can in principle rotate, the time scale for this is orders of magnitude longer than is viable. This instead implies that either the zygote always divides along the ZP’s long axis or that the ZP is soft at this stage and so able to remould itself around the two cells (25–27).

The next set of divisions, from two to four cells arranged in a tetrahedron, also poses interesting questions (50, 51). Our model predicts that cell division along the long axis (as in Hertwig’s rule) is most likely to lead to the observed tetrahedron shape (47). However, we also find there is a degree of stability to this: up to a 25^°^ variation about the long axis still leads to a tetrahedral arrangement. However, it is worth pointing out that this conclusion should be treated with care since the four-cell tetrahedron shape cannot be fully understood with our two-dimensional model.

After this stage, cells start to compact, with an increase in cell-to-cell contact area and associated substantial increase in the angle between cells (by 61^°^ in mouse and 77^°^ in human). Our model shows that this increase cannot be caused simply by increased adhesion between cells, but rather must involve a change in cortical tension around the cell. This could be just an increase in the unadhered tension constant Γ, but cannot be due to just a decrease in the adhered constant *γ*. However, in mouse, our simulation predicts the best fit to the experimental data if both Γ is doubled and *γ* reduced by 20% (see sketch in Fig. 4C) (22, 71).

Following compaction, a subset of cells internalise by moving to the centre of the developing morula and losing all contact with the outside. Since this involves cells splitting into apolar cells (which internalise) and polar cells (that remain on the outside), there are now four possible tension coefficients: Γ and *γ* for both apolar and polar cells (9, 35, 58). Our model shows that at least three of these must change to achieve complete internalisation, with the best result for mouse involving all four values changing. In particular, we predict that polar cells must decrease both Γ and *γ*, whereas apolar cells must increase both (see Fig. 5H). This difference in adhered tension coefficient *γ* has previously been noted (58), but has so far been overlooked as a facilitator of internalisation.

It is interesting to continue to follow the angle between cells during internalisation. We find that, if only the unadhered tension constant Γ is decreased, this would cause cells to start to decompact, with a decrease in cell-to-cell contact area. While small levels of decompaction followed by recompaction have been observed in both mouse and human, this is not common (59, 60). This suggests that the adhered tension constant *γ* must also increase during internalisation. Further, our model predicts that this should then lead to even greater compaction, with the angle between cells continuing to increase beyond the 146^°^ found in mouse and 158^°^ in human. Future experiments should be able to test this.

Both intra- and intercellular noise are an unavoidable feature of biological systems, often undesirable and occasionally beneficial. Here, we investigated two types of noise: “passive” noise with constant magnitude and “active” noise whose magnitude is linearly dependent on the local cortical tension *k*_ten_. In many cases we found noise to be detrimental, decreasing the compaction angle and reducing the level of internalisation. However, our simulations demonstrated that active noise can cause an increased angle between cells during compaction and that both types of noise are able to aid internalisation in cases that would otherwise fail, thereby conferring a degree of stability to the system. These results suggest that the role of noise may be to allow the same outcome (successful compaction and internalisation) with smaller changes to the actomyosin network than would otherwise be needed.

Later divisions are unlikely to be governed by the same rules as early cell cleavage, not least because cells start to acquire distinct identities and must be arranged in prescribed patterns. There has long been a debate on whether trophectoderm (TE) fate is decided by cell position or by polarisation (whether a cell contains an apical domain). Our model conclusively supports the latter by showing that even in the extreme, unrealistic case that the apical domain is inherited by the most internal daughter cell, the cells quickly switch position so that the cell on the outside (which will become TE) is the one with the apical domain.

The choice of cell division axis at these later stages seems to be more complex than at earlier times. Division is either “symmetric” (with cells dividing along their long axis) or “asymmetric” (with division such that the apical domain is inherited by only one daughter cell). It has been suggested that the choice of division axis is driven by the aspect ratio of the cell, with more elongated cells dividing symmetrically (67). This is exactly what our model shows: we predict that cells with an aspect ratio over two divide symmetrically, whereas more spherical cells divide asymmetrically. Further, as in Watanabe et al. (12), we find an interesting switch in the distribution of division angles from negative to positive skew between the 8-16 to the 16-32-cell stages.

Finally, we examined the similarities and differences between mouse and human. Our simulation works equally well in both cases, suggesting that the underlying biophysical mechanisms are conserved, potentially across many different mammals. However, although the fundamental mechanisms may be the same, the associated parameters are not. We predict that the pre-compaction tension coefficients (both Γ and *γ*) are 50% larger in human than in mouse. In addition, during compaction, these coefficients must change differently: we find the best fit to experimental data if the unadhered tension coefficient Γ increases (for both human and mouse), but the adhered coefficient *γ* increases in human and decreases in mouse.

To assess our results, it is important to be aware of the limitations of our modelling approach. First, our simulation is two-dimensional, despite the three-dimensional nature of the embryo. This is necessary to ensure that simulations run in reasonable time. However, we believe this is less of an issue than might initially be thought: the key questions that we address and that are generally of interest at this developmental stage are tractable with a two-dimensional model, negating the need for a more complex three-dimensional model.

Other model limitations include the functional form we choose for forces. For example, we assume the tension force to be governed by a Hooke-like law. Although there is likely to be a regime where this is accurate, it is probably more exact to treat the cortex as a viscoelastic material. Similarly, we choose a simple form for the curvature force, which in reality is likely to be much more complex.

We also ignore the internal structure of the cells. For example, we include the role of actin only via its contribution to the cortical tension and curvature, without any attempt to model the detailed spatiotemporal actomyosin dynamics. Further, we do not explicitly consider cell signalling, either within or between cells. Instead, we simply impose abrupt changes to the model parameters at key points of development such as compaction and internalisation.

Future work could address some of this simplifications by, for example, considering more complex forms of the forces or more realistic cell division (such as the observed blastomere “rounding up” just before cleavage (12, 72)). We have assumed that the cell preferred area *A*_0_ and length *L*_0_ are constants. It would be interesting to relax this assumption and allow these values to change dynamically. In addition, although membrane curvature plays a relatively small role in our model, more involved implementations (perhaps including spontaneous membrane curvature) could be investigated in future.

Our model focussed exclusively on the earliest stages of embryogenesis, up until the mature morula. However, it could easily be extended to study subsequent processes such as inner cell mass (ICM) differentiation into epiblast and hypoblast, blastocoel formation and ICM cell sorting, resulting in the blastocyst. These models could also include descriptions of relevant gene regulatory networks (GRNs), such as those involved in ICM and TE differentiation. Finally, with only minor changes, our model could be used to study other parts of development (such as neurulation), related organisms (such as cattle), and stem-cell organoids such as blastoids (29, 73–76).

Our model has led to a number of key predictions and potential novel understanding, not least concerning the underlying biophysical rules that may span mammalian development and perhaps even extend throughout other parts of the animal kingdom. Notwithstanding the obvious importance of biochemistry and cell signalling, we have demonstrated that the biophysical concepts that govern the earliest stages of development are remarkably simple and consist of only a handful of forces such as cortical tension and cell-to-cell adhesion. Future modelling is likely to continue to make progress on this important topic, with potential benefits to globally important applications such as *in vitro* fertilisation and conservation of endangered species.

## Methods

### Model simulation

We implement our model in C++ using a time step of Δ*t* = 0.1s and an initial vertex spacing of Δ*L* = 1*μ*m. At each time step, we calculate the total force on each vertex, convert this to a velocity and move the vertices the appropriate distance. After each time step, to ensure that vertices are always approximately Δ*L* apart, we remesh if necessary by removing or adding vertices when two vertices move too close together or too far apart. When cells divide, a single polygon is converted into two polygons, with new vertices added as required. Vertices that are within a fixed capture radius, *R*_capture_, become adhered to each other and start to feel the adhesion force. Adhered vertices can also unadhere if they move sufficiently far apart. In order to prevent cells from overlapping each other, we check for cell intersection at each time step and move any overlapping vertices either side of the cell intersection midpoint. The initial condition used depends on the precise question being investigated, but is either one, two, four or six cells. All code and related documents can be found at https://github.com/Cell-gorl/vertex_model_early_emb. For a video showing single run of the entire simulation form single fertilised egg up to the mature morula, see Supplementary Movie S1.

### Parameter values

Parameter values are either taken from the literature or inferred from experimental data as explained in the text. For the frictional coefficient we use *k*_fric_ = 33.3pNs*μ*m^−3^ (77–79). The volume force coefficient was chosen as *k*_vol_ = 3nN*μ*m^−2^ (9), the curvature force coefficient as *k*_curv_ = 0.1aNm (80) and the adhesion coefficient *k*_adh_ = 0.01pN*μ*m^−2^ (41, 42). The tension coefficient, *k*_ten_, and particularly its values at different parts of the cell, *γ* and Γ, are fit to experimental data as explained in the main text. For a full list of parameters and their values, see the table in the Supporting Information.

## Funding information

DMR gratefully acknowledges financial support from the Medical Research Council (MR/P022405/1), and from the Biotechnology and Biological Sciences Research Council and the National Centre for the Replacement, Refinement and Reduction of Animals in Research (NC/X002268/1). KTA acknowledges the financial support of the EPSRC via grant EP/T017856/1. PS thanks the National Centre for the Replacement, Refinement and Reduction of Animals in Research for funding this work via an Early Career Engagement award (NC/ECE0028/1). The funders had no role in study design, data collection and analysis, decision to publish, or preparation of the manuscript.

## Acknowledgements

We thank Melanie White for critically reading the manuscript.

## Supporting information

**1. Supporting Text S1** - further details on the model and its implementation

**2. Supporting Movie S2** - example of a complete simulation from a single fertilised egg to the mature morula

## Notes

The authors have no conflicts of interest to declare.

### Competing Interest Statement

The authors have declared no competing interest.

