## Supporting Text S1 for "How simple physics drives the earliest stages of embryogenesis"

#### 1 The vertex model

##### 1.1 Overall model framework

We represent the developing morula using a vertex model, with each cell represented by a polygon. Each polygon is defined by a set of vertices and straight edges that join pairs of adjacent vertices (see Fig. 1). Our primary focus is on the movement of the vertices, which captures how cells change position and shape over time.

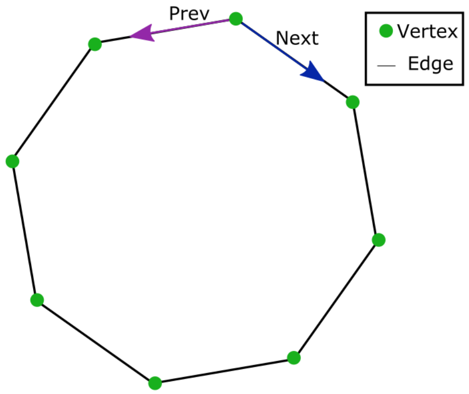


Figure 1: Sketch of how cells are represented in the vertex model framework.

Most vertex models of multicellular systems share vertices between neighbouring cells. For example, this approach is typically used to describe epithelial sheets where there can be no gaps between cells. However, here, we instead use an "unshared" vertex model, where each cell has its own set of vertices. Nearby vertices on distinct cells can still interact so that cells can adhere together, but cells are also free to separate from their neighbours.

We implement the model in C++ using a doubly linked list to describe each cell's set of vertices. This makes it easy and computationally quick to add and remove vertices when needed. Since our model is two-dimensional, each vertex position is described by its x- and y-coordinate.

#### 1.2 Forces on vertices

During the simulations we implement four biophysical forces on each vertex: cortical tension (stemming from the membrane and the associated underlying actin cortex), a volume force (due to approximate volume conservation of the cytoplasm), a curvature force (from the arrangement of phospholipids in the membrane and the mechanical properties of the actin cortex), and adhesion (between adjacent cells). We call the sum of these forces $F_{\text{total}}$.

There is a also a drag force on each vertex given by $F_{\text{drag}} = k_{\text{fric}} h_{\text{cell}} \Delta L v$ where $k_{\text{fric}}$ is the frictional coefficient and $v$ the vertex velocity. Here, $h_{\text{cell}} \Delta L$ can be thought of as the area of the little patch of membrane associated with the vertex.

In total this gives $F_{\text{total}} - F_{\text{drag}} = ma$ where $m$ is the mass of the little region of the cell associated with the vertex and $a$ is its acceleration. However, at the low Reynolds numbers that characterise biology at micrometre length scales, fluid dynamics is dominated by viscous effects and so, as is almost always assumed, we set $a = 0$ at all times. This leaves $F_{\text{drag}} = F_{\text{total}}$, which can be solved to find the velocity of each vertex.

#### 1.3 The radius of curvature

Finding the curvature force requires calculating the radius of curvature at each vertex. For a given vertex $v$, we do this by first finding the two neighbouring vertices to $v$, one on each side. Rather than choosing the adjacent vertices, we instead find the vertices that are as close as possible to a distance of $\lambda$ from $v$. This is important to ensure that the simulation is independent of the lattice spacing $\Delta L$. We normally choose $\lambda = 0.95\mu$m. The radius of curvature is then found as the radius of the circle that passes through $v$ and the two vertices either side.

#### 1.4 Adhered vertices

The adhesion force is different to the other three forces in that it only operates between pairs of vertices (on different cells) that are sufficiently close to each other. We refer to pairs of vertices that feel the adhesion force as adhered vertices and keep track of which vertices are adhered.

A pair of vertices become adhered if they move within a the capture radius $R_{\text{capture}}$ of each other. They then remain adhered until they move apart by more than the release radius $R_{\text{release}}$. For simplicity we always assume that $R_{\text{capture}} = R_{\text{release}}$ and typically use a value of $R_{\text{capture}} = 0.6\mu$m.

#### 1.5 The apical domain

During later divisions, we often need to define the apical domain for each cell. To do this, we first identify the centre of the morula, *i.e.* the centre of mass of all the cells. For a given cell, we then find the unadhered vertex that is furthest from the centre of the morula. We then find the set of unadhered vertices that contain this vertex and are all connected to each other. This set defines a region of the cell membrane. If this set has length longer than a fifth of the total cell perimeter, then the entire region is labelled as the apical domain.

#### 1.6 Remeshing

We start our simulation with vertices that are distance $\Delta L$ from each other. However, as the simulation proceeds the distance between neighbouring vertices can increase and decrease. To maintain an approximate distance $\Delta L$ between vertices, we remesh whenever vertices move too far apart from or too near to each other (see Fig. 2).

To do this we periodically check (after each time step) the distance between neighbouring vertices. If this distance is every more than $2\Delta L$, then we insert as many new vertices as possible between the two vertices to ensure that the spacing is as close to $\Delta L$ as possible. New vertices are evenly spaced and are inserted along the straight line connecting the two overly-separated vertices. Conversely, if the distance between two neighbouring vertices is every less than $0.5\Delta L$, one of the vertices is removed.

In the case of adding vertices, if either of the two overly-separated vertices are adhered, we also adhere the new vertices to associated new vertices on the adhered cell. Similarly, if one of the two vertices is part of the apical domain, the new vertices are also part of the apical domain.

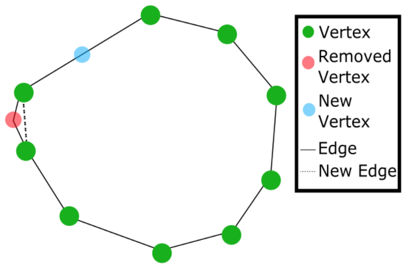


Figure 2: Schematic of how the remesh function works

#### 1.7 The zona pellucida

In our simulation the zona pellucida (ZP) is a fixed region within which all vertices must lie. If a vertex tries to move outside this region then it is prevented from doing so and returned to its previous position. Altohugh more general shapes would be easy to implement, we always assume the ZP is elliptical.

#### 1.8 Preventing intersection between cells

Unless care is taken vertices can try to move within the interior of cells. This is an intersection issue and occurs because a give vertex has tried to cross an edge between another pair of vertices. This can happen either within the same cell (*i.e.* self-intersection) or between two different cells (where one cell tries to cross into another cell).

To fix this we implement a non-intersection algorithm that runs after each time step. If any intersection is found, the offending vertices are partially moved back to their previous position so that there is no longer any intersection.

#### 1.9 Cell division

The first stage of a cell division involves identifying the axis of division. We consider various options, including division along the shortest axis, the longest axis or a randomly chosen axis. We also sometimes allow the position of the apical domain to control the division axis. In either case, the division axis is defined by two vertices on opposite sides of the cell.

To divide the cells, the vertices are split into two groups and each identified with a new cell. The resting area $A_0$ of each new cell is set as half that of the mother cell. The remesh function is then called on both daughter cells in order to ensure that the line of division is evenly covered by new vertices.

The timings of cell division are chosen stochastically from a (truncated) normal distribution with given mean $\tau$ and standard deviation $\sigma_\tau$. Although unlikely, we prevent cells from dividing earlier than $\tau - 3\sigma_\tau$.

### 2 Extraction of quantities from the simulations

#### 2.1 The angle between cells, $\theta$

The average angle $\theta$ between two adhered cells is an important quantity that has been measured experimentally in both mouse and human. To extract $\theta$ from our simulations, we first identify the pair of vertices at the end of the adhered region, one on each cell (see Fig. 3). For each of these vertices, we consider the next and previous vertex in that cell, calculating the angle between them in each case and ensuring each angle is smaller than 180°. We also calculate the same angle by jumping more than one vertex forward, taking the final angle as the largest of all the angles. This gives two angles, $\theta_1$ and $\theta_2$, one for each end vertex. The angle between the cells is then given by $\theta = 360° - \theta_1 - \theta_2$. Averaging over many simulations provides the final average angle between cells that can be compared to experiment.

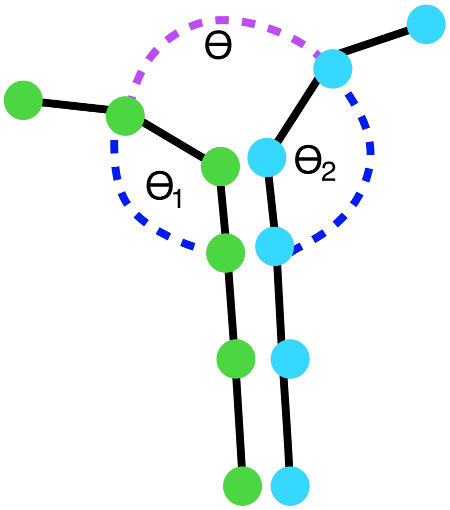


Figure 3: Schematic of the angle $\theta$ between two cells, which we calculate from the two angles $\theta_1$ and $\theta_2$.

#### 2.2 The compaction parameter, $\alpha$

The other relevant experimental value is the compaction parameter, defined as $\alpha = \gamma_{cc}/2\gamma_{cm}$, where $\gamma_{cc}$ is the surface tension at a cell-to-cell interface and $\gamma_{cm}$ is the surface tension at a cell-medium interface. This is harder to measure in our simulation, largely because of the difficulty in translating the distinct biophysical forces that we consider into the associated surface tension. Surface tension is often understood as being related to two facets: an inward force due to the imbalance of molecular attractive forces at the surface and a tangential force acting on the surface that arises due to the decreased pressure parallel to the surface [1].

In our simulations we assume that surface tension is caused by the resultant normal force acting at a given part of the cell membrane. The experiments that we compare to were performed with micropipette aspiration and so are unlikely to capture only the effect of cortical tension [2]. Thus we include all of our forces in calculating the resultant normal force: cortical tension, curvature, volume conservation and adhesion.

The surface tension at a cell-to-cell interface, $\gamma_{cc}$, is found as the resultant normal force averaged over all vertices that are part of the adhered region, *i.e.* all vertices bound to the neighbouring cell. Conversely, the surface tension at the cell-medium interface, $\gamma_{cm}$, is found by averaging the normal force over all unbound vertices, *i.e.* those that are in contact with the medium.

The compaction parameter $\alpha$ is then found as the ratio of the two surface tensions, as given by the expression above. Even if our calculation is not a perfect representation of the surface tension, the fact that only the ratio of surface tensions is needed means that our value of $\alpha$ is still likely to be a useful estimate of the experimentally-measured value [2].

### 3 Comparison of cell numbers in two and three dimensions

Our model is two-dimensional, but represents the full three-dimensional developing morula. Thus in order to study $N$ 3D cells, we need to know the equivalent number of 2D cells in our simulation.

To determine this, we used a 3D cellular Potts model (CPM) of the morula that we have developed separately to this work. We allowed the CPM to run until the $N$-cell stage and then considered a 2D slice through the centre of the morula.

For a spherical cell of radius $R$, we expect its volume to be $V = \frac{4}{3}\pi R^3$ and its maximum cross-sectional area to be $A_{\max} = \pi R^2$. Typically only a fraction of this maximum area will be seen in each slice. Within our 2D slice we count a cell as present/visible if its area is greater than $0.5A_{\max}$.

By running our CPM many times, we are able to find a map between the number of 3D cells, $N$, and the average number of cells within a slice. We then use the average number of cells in 2D (rounded to the nearest integer) within our vertex model simulation. Our results are shown in Table 1.

| Number of cells in 3D | Av. number of cells in 2D | St. dev. of number of cells in 2D |
|---|---|---|
| 4 | 2.94 | 0.79 |
| 8 | 4.72 | 0.80 |
| 16 | 7.13 | 1.16 |
| 32 | 11.20 | 1.35 |

Table 1: Map between number of cells in 3D and number of cells in our 2D simulation

### 4 Model and simulation parameters

In Table 2 we summarise all model and simulation parameters along with their values (or range of values).

| Parameter | Symbol | Value | Reference |
|---|---|---|---|
| Tension coefficient | $k_{\text{ten}}$ | 0.01-100 $\text{pN}\mu\text{m}^{-1}$ | [2] |
| Volume coefficient | $k_{\text{vol}}$ | 3 $\text{nN}\mu\text{m}^{-2}$ | [3] |
| Curvature coefficient | $k_{\text{curv}}$ | 0.1 $\text{aNm}^{-1}$ | [4] |
| Adhesion coefficient | $k_{\text{adh}}$ | 0.01 $\text{pN}\mu\text{m}^{-2}$ | [5, 6] |
| Frictional coefficient | $k_{\text{fric}}$ | 33.3 $\text{pNs}\mu\text{m}^{-3}$ | [7, 8, 9] |
| Spontaneous curvature | $H_0$ | 0 $\mu\text{m}^{-1}$ | - |
| Initial cell radius | $R_{\text{cell}}$ | 35-40 $\mu$m | [10, 11] |
| ZP radius | $R_{\text{ZP}}$ | 40-45 $\mu$m | [12, 13] |
| Capture radius | $R_{\text{capture}}$ | 0.6 $\mu$m | - |
| Cell height | $h_{\text{cell}}$ | 70 $\mu$m | [10, 11] |
| Noise magnitude | $\zeta$ | 0.001 $\text{N/s}^{1/2}$ | - |
| Mean division time | $\tau$ | 18-20 ($1^{\text{st}}$-$2^{\text{nd}}$), 12-14 ($3^{\text{rd}}$-$5^{\text{th}}$) hrs | [14] |
| Division time st. dev. | $\sigma_\tau$ | 1 ($1^{\text{st}}$-$2^{\text{nd}}$), 2 ($3^{\text{rd}}$-$5^{\text{th}}$) hrs | [15] |
| Vertex spacing | $\Delta L$ | 1 $\mu$m | - |
| Time step | $\Delta t$ | 0.1 s | - |
| Curvature jump | $\lambda$ | 0.95 $\mu$m | - |

Table 2: Model and simulation parameters
